# Zinc preserves translational proteostasis to counteract aging and proteotoxicity in *Caenorhabditis elegans*

**DOI:** 10.64898/2026.10.07.757281

**Authors:** Arun Kumar, Kamila P. Liput, Katarzyna Jonak, Rafał Mazgaj, Spoorthy Gowda, Maja Miążek, Kevin J. Waldron, Ulrike Topf

## Abstract

The progressive collapse of protein homeostasis is a defining feature of aging, yet upstream events deteriorating protein synthesis remain poorly defined. Zinc is an essential micronutrient and structural cofactor, but its role in the aging ribosomal proteome is poorly understood. Here, using *Caenorhabditis elegans*, we show that the whole-body zinc-to-sulfur ratio declines by ∼90% between day 1 and day 13 of adulthood. Supplementing aged animals with a low dose of ZnCl_2_ (10 μM) restored organismal zinc levels and improved healthspan, thermotolerance, and oxidative stress resistance, whereas higher doses were toxic. Quantitative proteomics revealed that structural constituents of the ribosome were the most enriched functional category altered by zinc, accompanied by robust induction of metallothionein MTL-1. Zinc-supplemented animals exhibited increased protein synthesis and a transient increase in ribosomal proteins, while 45S pre-rRNA levels remained unaffected, indicating that zinc acts on ribosome structure rather than biogenesis. Zinc supplementation coordinately reversed the age-associated decline of antioxidant, chaperone, proteasome, and energy-metabolism factors. Furthermore, zinc conferred protection in two proteotoxicity models, suppressing polyglutamine aggregation and delaying amyloid-beta-induced paralysis. Our findings establish that zinc preserves translational proteostasis, identifying late-life zinc supplementation as a translatable micronutrient intervention against age-related proteinopathies.

## Introduction

Aging is characterized by a progressive loss of physiological integrity that leads to impaired cellular function and ends in organismal death. It constitutes the primary risk factor for major chronic diseases, including cancer, cardiovascular disease, and neurodegeneration [1]. Among the interconnected aging hallmarks that underlie this decline, the loss of protein homeostasis (proteostasis), characterized by the collapse of the integrated network of molecular chaperones, the ubiquitin-proteasome system, and autophagy that maintains the cellular proteome in a soluble, functional state, occupies a central position [1, 2]. In the nematode *Caenorhabditis elegans*, proteostasis capacity does not decline gradually across the entire lifespan; rather, it collapses abruptly at the onset of adulthood, so that the ability to buffer aggregation-prone proteins is already compromised in early post-reproductive animals [3]. Collapse in cellular proteostasis is not merely permissive for disease. Endogenous proteins can become detergent-insoluble and aggregation-prone with age even in the absence of any disease-associated transgene, and this inherent aggregation load aggravates the toxicity of disease proteins [4]. Understanding what triggers the deterioration of proteostasis, and whether it can be reversed in animals that are already old, is therefore a question of considerable biomedical importance.

An understudied aspect of proteostatic decline concerns the fidelity and output of the protein synthesis machinery itself. The ribosome is an essential part of the proteostasis network. The accuracy with which it decodes mRNA sets the quality of every protein a cell makes, and errors in decoding generate misfolded species that must subsequently be handled by downstream quality-control systems [5, 6]. Both, the overall rate of translation and the fidelity of decoding change with age, and these two parameters have opposing relation to longevity. Genetic or pharmacological reduction of bulk translation extends lifespan, whereas a loss of decoding accuracy is associated with aged and diseased phenotypes [7, 8]. Strikingly, increasing translational fidelity is sufficient to extend lifespan. An error-restrictive mutation in the small subunit (SSU) ribosomal protein RPS23/uS12 that stabilizes the decoding center prolongs the life of yeast, flies, and worms, and the anti-aging drugs rapamycin, torin1, and trametinib act at least in part by enhancing decoding accuracy [9, 10]. Conversely, the aminoglycoside paromomycin, which binds the SSU decoding site and relaxes the stringency of codon-anticodon proofreading, increases mistranslation and proteotoxic stress and is widely used as a pharmacological tool to provoke error-prone translation [9, 11, 12]. That translational fidelity coevolves with species longevity further underscores the accuracy of protein synthesis as a determinant of organismal healthspan [13]. Despite this, the biochemical cofactors that maintain ribosome integrity during aging have received little attention.

Zinc is one such cofactor, and an unusually prevalent one. It is the second most abundant trace metal in living systems and an essential structural and catalytic component of many proteins. A large fraction of the eukaryotic proteome is predicted to bind zinc, much of it through cysteine-rich zinc-finger motifs that fold only when the metal is coordinated [14]. The ribosome is among the most zinc-dependent machines in the cell. Multiple ribosomal proteins carry C4-type zinc-finger motifs whose four cysteine residues coordinate a single structural zinc ion. In budding yeast, nine ribosomal proteins possess such motifs, however only eight of them have been crystallised with a bound zinc ion. Mutation of the zinc-coordinating cysteine residues of the ribosomal protein Rpl37a/eL37 in yeast abolishes metal binding, and one of the cysteine residues was essential for cell viability [15]. Similar findings were obtained for the yeast ribosomal protein Rps26/eS26 [16]. Zinc-binding ribosomal proteins are well conserved. However, few ribosomal proteins have been shown to bind zinc ions in *C. elegans* [17]. Nevertheless, the conserved C4-type motif and structural fold imply the metal binding, rendering the ribosome a substantial and structurally committed pool of cellular zinc.

Zinc homeostasis deteriorates in the elderly. Dietary zinc intake decreases when intestinal absorption becomes less efficient resulting in zinc deficiency. At least a quarter of the world’s population is at risk, with the aged disproportionately affected [18, 19]. Zinc deficiency in older people is linked to immune dysfunction, chronic inflammation, and cognitive impairment [19–21]. Consequently, zinc has attracted interest as a micronutrient intervention. Yet the field has focused primarily on zinc ions’ roles in immunity, redox balance, and transcription-factor function, while zinc ion depletion during aging and its consequences for translation and proteostasis has been less explored [20, 21]. We reasoned that the ribosome is a zinc-dependent machine, therefore a decline of cellular zinc ions with age might compromise ribosome integrity and translational proteostasis, and that restoring zinc ion levels to aged animals might reinforce the proteostasis network. We tested this hypothesis in *C. elegans*, an animal in which proteostatic collapse, protein-aggregation disease, and the metal composition of the whole organism can all be interrogated together. We deliberately supplemented animals only after they had reached the post-reproductive stage, so that any phenotype would reflect an effect on aging itself rather than on development, growth, or reproduction. We find that whole-body zinc ion levels decline steeply in post-reproductive animals. Replenishing it improved locomotion, stress resistance, and survival in a zinc-dependent manner. The dominant molecular signature of zinc supplementation is an enrichment of ribosomal proteins accompanied by increased protein synthesis, and zinc ion supplementation coordinately restores the antioxidant, chaperone, proteasome, and bioenergetic arms of the proteostasis network. Finally, zinc ion supplementation protected against both polyglutamine and amyloid-beta proteotoxicity by priming a selective, rather than bulk, autophagy program. Together, these results define a zinc-ribosome axis that links a common micronutrient deficit of old age to the integrity of protein synthesis, and suggest late-life zinc repletion as a mechanistically distinct intervention against the proteinopathies of aging.

## Results

### Trace-metal homeostasis is remodelled during adulthood, with a pronounced late-life collapse of zinc ions

To determine whether the bodily metal economy changes as animals age, we quantified six biologically important elements, zinc, manganese, iron, copper, calcium, and magnesium across the adult lifespan of wild-type (N2) animals by inductively coupled plasma optical emission spectrometry (ICP-OES), normalizing each metal to sulfur to control for biomass (**Fig. 1A-F**). Zinc exhibited the most pronounced age-dependent change of all the elements examined. The zinc-to-sulfur ratio increased modestly from day 1 to a peak at day 4 of adulthood (a 24% increase) and then declined, falling by 33% by day 7, by 60% by day 10, and by ∼90% by day 13, where it stabilized through day 16 (**Fig. 1A**; *p* < 0.001 for all time points versus day 1). The other elements followed distinct trajectories: manganese peaked at day 7 before returning toward baseline (**Fig. 1B**); iron and calcium increased in later life (**Fig. 1C and D**); copper increased early and then remained comparable to day 1 (**Fig. 1E**); and magnesium remained comparatively stable (**Fig. 1F**). The steep and sustained loss of zinc ions during the post-reproductive period identified zinc as the metal most evidently depleted by aging. This motivated us to ask whether restoring zinc levels could influence the aging process.

**Figure 1.**
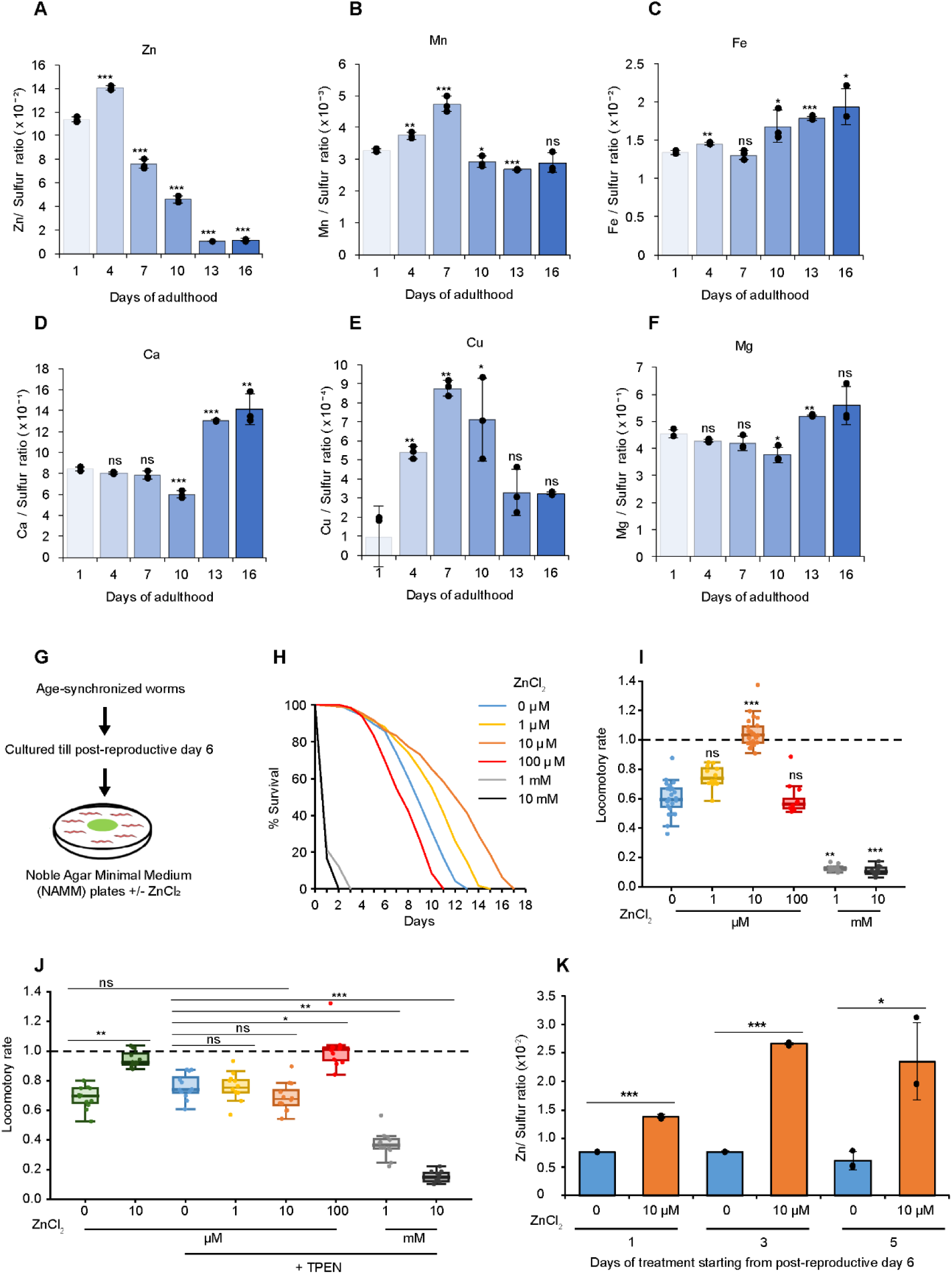
Whole-body zinc declines steeply in post-reproductive animals, and late-life zinc supplementation extends survival and improves locomotion in a zinc-dependent manner. (A-F) Age-dependent trace-metal content of wild-type (N2) animals across adulthood (days 1, 4, 7, 10, 13, 16), measured by ICP-OES and expressed as the metal-to-sulfur ratio: (A) zinc, (B) manganese, (C) iron, (D) calcium, (E) copper, (F) magnesium. Data are mean ± SD, *n* = 3 biological replicates; individual replicates shown as points. Significance versus day 1 by unpaired two-tailed *t*-test; \**p* < 0.05, \*\**p* < 0.01, \*\*\**p* < 0.001, ns not significant. (G) Experimental scheme: age-synchronized animals were cultured to post-reproductive day 6 and transferred to Noble Agar Minimal Medium (NAMM) supplemented with ZnCl_2_. (H) Kaplan-Meier survival of post-reproductive N2 animals on the indicated ZnCl_2_ concentrations (0, 1, 10, 100 μM, 1, 10 mM); *n* = 300 per condition of three biological replicates. Median survival was maximal at 10 μM (12 days) and collapsed to 1 day at ≥ 1 mM. (I) Normalized locomotory rate over 72 h (WMicrotracker) as a function of ZnCl_2_ dose; box plots show well-level values. 10 μM significantly increased locomotion versus 0 μM (Kruskal-Wallis with Dunn’s post-hoc, BH-FDR; *p* < 0.001); 1 mM and 10 mM significantly suppressed it (*p* < 0.01, *p* < 0.001); 1 and 100 μM were not significant. (J) Locomotory rate for ZnCl_2_ alone versus ZnCl_2_ co-administered with the metal chelator TPEN (200 μM). The 10 μM benefit was abolished by TPEN (10 μM + TPEN not significant versus 0 μM control). (K) ICP-OES zinc-to-sulfur ratio in animals treated with 0 or 10 μM ZnCl_2_ for 1, 3, or 5 days; mean ± SD, *n* = 3; *p < 0.001* at days 1 and 3 and *p* < 0.05 at day 5 versus matched untreated control. WT, wild type.

### Low-dose zinc supplementation of aged animals extends survival and improves locomotion

To test the consequences of replenishing zinc after its natural decline, we established a late-life supplementation paradigm. Age-synchronized animals were cultured under standard conditions until post-reproductive day 6, which is beyond the developmental and reproductive phases, and then transferred to Noble Agar Minimal Medium (NAMM), a defined low-background substrate, supplemented with ZnCl_2_ across a wide concentration range (**Fig. 1G**). Supplementation began only in aged adults, and thus, any resulting phenotype reflects an action on aging *per se* rather than on growth or fecundity.

Zinc influenced survival in a biphasic, dose-dependent manner (**Fig. 1H**). Relative to control (0 μM), 10 μM ZnCl_2_ conferred the greatest benefit, extending median survival from 9 to 12 days (replicate-median 13.0 days), with 1 μM producing an intermediate extension to 11 days. In contrast, millimolar concentrations were acutely toxic: at 1 mM and 10 mM, median survival collapsed to a single day, defining a narrow therapeutic window with a sharp toxicity threshold. Feeding dead bacteria while supplementing worms with zinc resulted in a similar effect on longevity and thus was independent from the active bacterial metabolism (**Supplementary Fig. 1A**). A parallel dose-response of neuromuscular function, measured as bulk locomotory activity over 72 h in liquid culture, mirrored the survival data (**Fig. 1I and Supplementary Fig. 1B**). Locomotion was maximal at 10 μM, which raised the normalized locomotory rate significantly above the untreated control (median 1.03 versus 0.60; Kruskal-Wallis with Dunn’s post-hoc, *p* < 0.001), whereas 1 μM and 100 μM were statistically indistinguishable from control, and 1 mM and 10 mM strongly suppressed movement (*p* < 0.01 and *p* < 0.001, respectively). We therefore adopted 10 μM ZnCl_2_ as the optimal dose for all subsequent mechanistic experiments. To assess whether the zinc-associated locomotory effect was retained at the level of individual worm movement, 50-60 individual worms were recorded and tracked with WormLab per condition. Time based crawling analysis was performed at post-reproductive day 6 worms (day 0) and worms were then maintained on NAMM supplemented with or without 10 μM ZnCl_2_ and re-recorded at day 1 (24 h) and day 3 (72 h) of treatment. Histograms of crawling speed revealed a progressive leftward shift (decrease in positive speed values, positive values indicating dominant forward motion) in the speed distribution from day 0 to day 3, consistent with age-related decline in post-reproductive animals (**Supplementary Fig. 1D**). Zinc supplementation partially counteracted this trend by day 3, the speed distribution in zinc-treated worms shifted rightward (toward higher positive values), suggesting an improved neuromuscular capacity. Supplemented and non-supplemented worms did show qualitative differences in the range of movement; lack of zinc impeded the individual’s ability to explore the new area compared to zinc supplementation as shown in representative images from three biological replicates (**Supplementary Figure 1E)**. The worms still displayed sinusoidal locomotory pattern for all the conditions and time points (**Supplementary Figure 1F**). This trend corroborates the direction of the zinc effect at single-worm resolution, complementing the population-level, dose-resolved result from the liquid-phase assay.

To establish that the locomotor benefit was caused by bioavailable zinc we co-administered the high-affinity, membrane-permeant metal chelator TPEN at a concentration of 200 μM (**Fig. 1J and Supplementary Fig. 1C**). Whereas 10 μM zinc alone significantly increased locomotion relative to control (median 0.92; *p* < 0.01), the 10 μM plus TPEN condition was statistically indistinguishable from untreated animals (median 0.68; not significant). The TPEN-sensitivity of the phenotype demonstrates that zinc availability, rather than any off-target effect, underlies the improvement in neuromuscular function.

Finally, we confirmed that our supplementation regimen reloads the aged animal with zinc. ICP-OES of animals treated with 10 μM ZnCl_2_ for 1, 3, or 5 days showed that supplementation increased the zinc-to-sulfur ratio at every time point compared to non-supplemented condition, (**Fig. 1K**; *p < 0.001* at days 1 and 3*, p < 0.05* at day 5). Thus, exogenous zinc is taken up by post-reproductive animals and reverses the age-associated depletion documented in Fig. 1A. Together, these data show that low-dose zinc supplementation of aged animals extends survival and improves healthspan, and that it is accompanied by restoration of the whole-organism zinc levels.

### Quantitative proteomics identifies the ribosome as a target of zinc supplementation

To understand why zinc improves the physiology of aged animals in an unbiased manner, we performed tandem mass tag (TMT) quantitative proteomics on animals treated with 0 or 10 μM ZnCl_2_, comparing the two conditions at day 1 and day 3 of treatment (**Supplementary Table 1**). After 24 hours, zinc supplementation reshaped the aged proteome substantially, with 236 proteins significantly upregulated and 249 proteins significantly downregulated (**Fig. 2A and Supplementary Fig. 2A**). Proteomic changes subsided by 72 hours, suggesting an acute adaptive homeostatic response rather than a sustained chronic shift in the aging proteome (**Supplementary Fig. 2B**). Prominent among the induced proteins was metallothionein-1 (MTL-1), the canonical zinc-buffering protein and a reporter of bioavailable cellular zinc, which was strongly upregulated by supplementation (**Fig. 2A**) and remained in the top upregulated proteins also after 72 hours of treatment (**Supplementary Fig. 2B**). We independently validated this response by RT-qPCR: *mtl-1* transcript was essentially unchanged during aging alone (day 0 versus untreated day 1) but was induced ∼4.8-fold by zinc at day 1 (**Fig. 2B**; *p* < 0.001 for zinc-treated versus both baseline and untreated day 1). The robust, zinc-selective induction of *mtl-1* confirms that supplemented animals activate a *bona fide* zinc-responsive transcriptional program and provides a molecular anchor for the supplementation effect. We next asked which functional categories dominate the zinc-responsive proteome. Gene Ontology enrichment analysis of the upregulated proteins (**Supplementary Table 2**) returned "structural constituent of the ribosome" as by far the most significantly and strongly enriched term (FDR = 9.2 x 10^-21^), well ahead of related categories such as structural molecule activity, unfolded protein binding, RNA binding, and protein-folding chaperone activity (**Fig. 2C**). The reciprocal analysis of downregulated proteins was enriched for translation factor and translation initiation factor activity together with diverse hydrolase and nucleotide-binding functions (**Fig. 2D**). This coherent signature, accumulation of ribosomal structural proteins alongside remodelling of translation factors, was substantiated by STRING network clustering, in which the ribosome emerged as the single largest and most connected cluster of proteins upregulated by zinc, mirrored by a translation cluster among the downregulated set (**Supplementary Fig. 2C, D**). Across two analytic frameworks (GO, **Fig. 2C, D**; STRING, **Supplementary Fig. 2C, D**), the ribosomal proteome was the outstanding molecular feature of the response to zinc supplementation, suggesting the ribosome as an important target through which zinc acts in aged animals.

**Figure 2.**
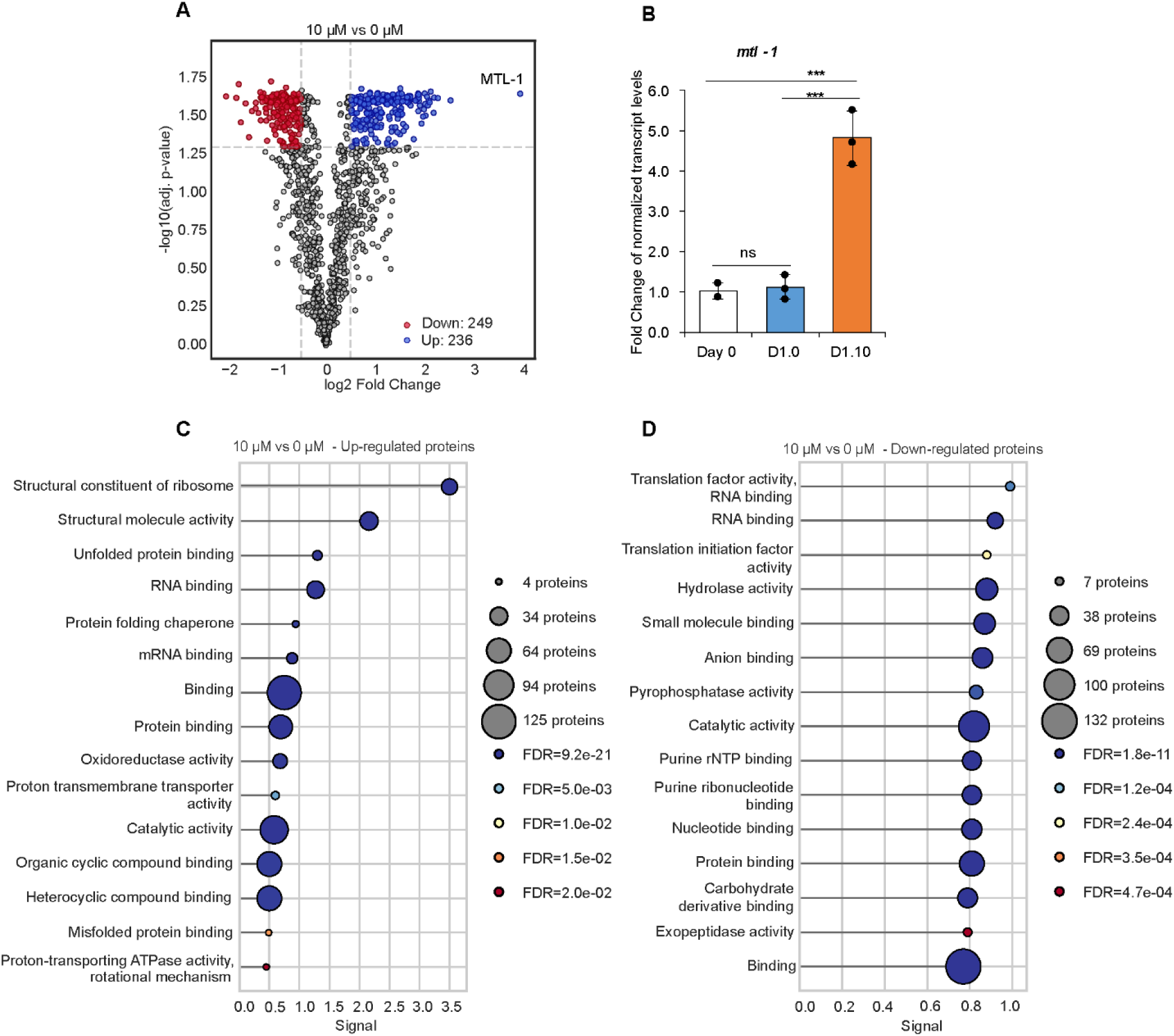
Quantitative proteomics identifies the ribosome as the dominant target of zinc supplementation. (A) Volcano plot of the aged proteome comparing 10 μM versus 0 μM ZnCl_2_ after one day of treatment. 236 proteins were significantly upregulated (blue) and 249 downregulated (red) at permutation-based FDR < 0.05 and log2 fold change ±0.5. Metallothionein-1 (MTL-1) is highlighted among the strongly induced proteins. (B) RT-qPCR validation of *mtl-1*; transcript is unchanged during aging alone (day 0 versus untreated day 1, ns) but induced ∼4.8-fold by zinc (D1.10), mean ± SD, *n* = 3; *p* < 0.001. (C) Gene Ontology enrichment analysis for Molecular Function (GOMF) of proteins upregulated upon addition of ZnCl_2_ at day 1. Top15 terms with the strongest signal according to STRING are presented. (D) GOMF enrichment of proteins downregulated upon addition of ZnCl_2_ at day 1. Top15 terms with the strongest signal according to STRING are presented.

### Zinc supplementation supports ribosome function during aging

An accumulation of ribosomal structural proteins does not, by itself, establish that translation is enhanced. We therefore examined the ribosomal signature more closely and tested translational output directly. Nine ribosomal proteins in *C. elegans* contain a CxxC motif, out of which five were proven to bind a zinc ion [17]. Our proteomic analysis identified six CxxC-motif-containing ribosomal proteins, two being significantly increased upon supplementation (RPS-26/eS26, RPS-27A/eS31) and two other showing tendency to increase (RPL-37A/eL43, RPS-12/eS12). However, the response was not restricted to zinc-binding ribosomal proteins. Interestingly, RPS-23, a ribosomal protein previously shown to be involved in positive regulation of translation fidelity during aging, was also significantly increased [9] (**Fig. 3A**). By RT-qPCR, we confirmed an increase in the two most upregulated ribosomal proteins, not zinc-binding *rpl-4* and zinc-binding *rps-26,* upon supplementation compared to non-supplemented worms at day 1 of treatment (**Fig. 3B and C**).

**Figure 3.**
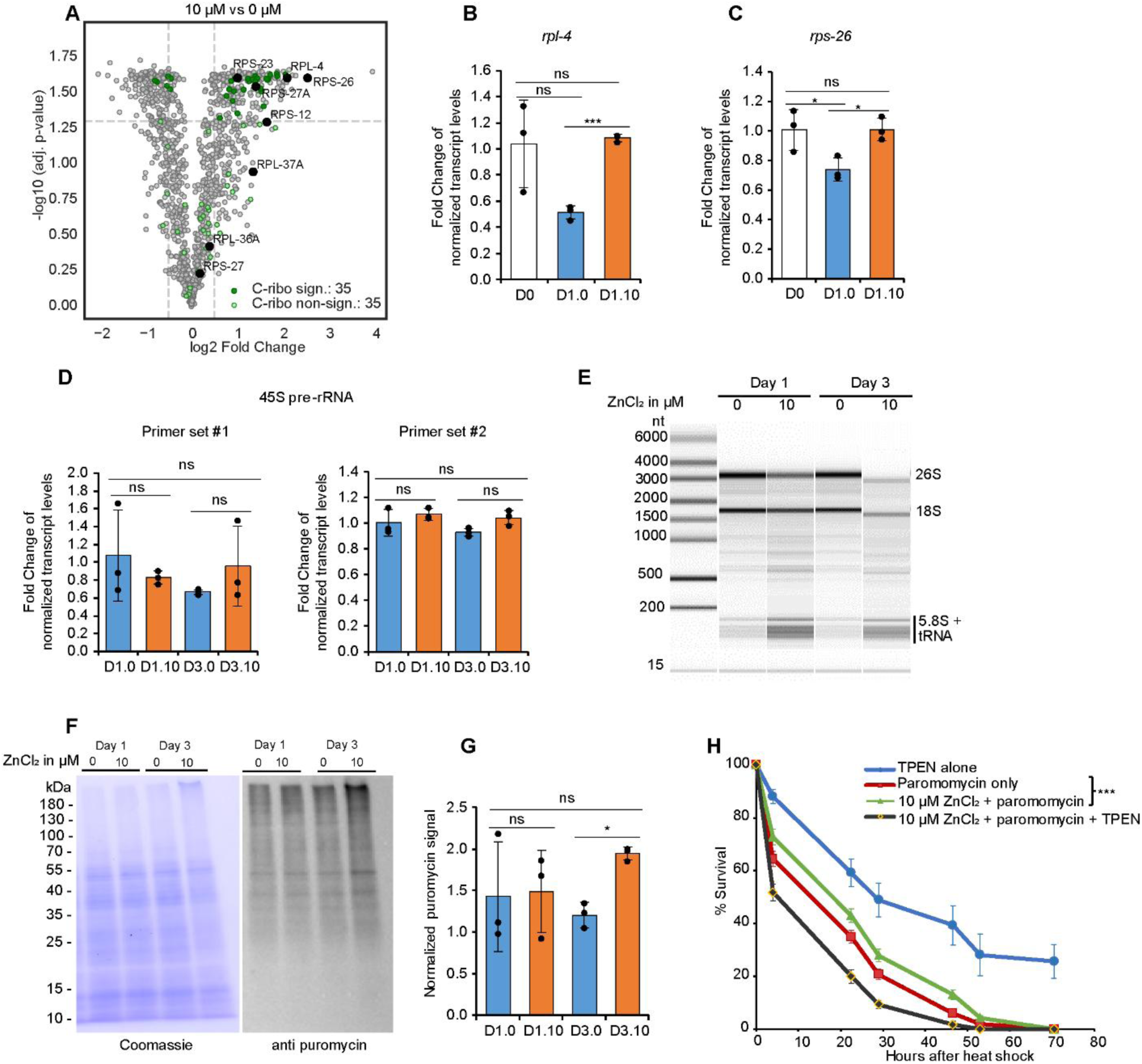
Zinc supplementation increases protein synthesis capacity. (A) Volcano plot (10 μM versus 0 μM, day 1) with the annotated cytoplasmic-ribosomal proteins overlaid (significant, dark green; non-significant, light green; permutation-based FDR < 0.05 and log2 fold change ±0.5). (B, C) RT-qPCR of *rpl-4* and *rps-26*: both decline with aging and are restored by zinc (mean +/- SD, *n* = 3; \**p* < 0.05, \*\*\**p* < 0.001, ns). (D) RT-qPCR using independent primer pairs (set #1 and #2) to amplify fragments of 45S pre-rRNA species. (E) Representative capillary-electrophoresis (Femto Pulse) traces of equal concentration of total RNA isolated from control animals (0 µM ZnCl_2_) or supplemented with 10 μM ZnCl_2_ for 1 day or 3 days. (F) SUnSET assay: representative Coomassie (loading) and anti-puromycin blots for 0 and 10 μM ZnCl_2_ at day 1 and day 3. (G) Quantification of puromycin vs. Coomassie signal; protein synthesis is significantly increased upon supplementation at day 3 (∼136% of control; *p* < 0.01), mean +/- SD, *n* = 3. (H) Post-heat-shock survival (35°C, 3 h) of aged animals under TPEN (200 μM) alone, paromomycin (0.5 mM) alone, 10 μM ZnCl_2_ + paromomycin (0.5 mM), or 10 μM ZnCl2 + paromomycin (0.5 mM) + TPEN (200 μM). Zinc improved survival over paromomycin alone (log-rank *p* < 0.001), and the benefit was reversed by TPEN (all pairwise comparisons *p* < 0.001). Mean +/- SD, *n* = 3.

To distinguish an effect on ribosomal protein abundance from a more general effect on ribosome biogenesis, we assessed abundance of ribosomal RNA. The levels of 45S pre-rRNA measured by RT-qPCR using two independent sets of primers revealed no change in the levels of this unprocessed precursor between zinc-treated and untreated animals at day 1 or day 3 (**Fig. 3D**). To examine the abundance of mature rRNA species we resolved equal concentrations of total RNA by capillary electrophoresis (**Fig. 3E**). Interestingly, zinc-supplemented animals showed a decrease in the 26S and 18S rRNA. The overall integrity of the rRNA was comparable between all samples. Thus, our data suggest that zinc does not act by accelerating ribosome biogenesis in general, but rather by restoring the composition and function of ribosomal proteins.

We then measured translational output directly using the SUnSET (surface sensing of translation) assay, which quantifies the incorporation of puromycin into nascent polypeptides [22]. Zinc-supplemented animals synthesized more protein, and the effect strengthened with the duration of treatment. Normalized puromycin incorporation was comparable between conditions at day 1 but was significantly elevated by zinc at day 3, reaching ∼136% of the untreated control (**Fig. 3F and G**; *p* < 0.01, 10 μM versus 0 μM at day 3). To determine whether this global translational benefit is functionally coupled to fidelity and organismal survival, we exploited paromomycin, which relaxes ribosomal decoding stringency to induce error-prone translation and proteotoxic stress [9]. Aged animals were subjected to a lethal proteotoxic heat shock in the presence of paromomycin, with or without zinc supplementation, and with or without the metal chelator TPEN (**Fig. 3H**). Paromomycin alone sharply reduced post-heat-shock survival; the addition of 10 μM zinc significantly improved survival at every timepoint (e.g., 43.0% versus 35.0% at 22.5 h; log-rank *p* < 0.001), consistent with zinc buffering the proteotoxic consequences of error-prone translation. Crucially, the protective effect of zinc was reversed when TPEN was included. TPEN-plus-zinc-plus-paromomycin animals died fastest of all groups (log-rank *p* < 0.001 versus every other condition), demonstrating that the survival benefit requires bioavailable intracellular zinc acting on the translation machinery. These data link zinc-dependent translational proteostasis to organismal survival under proteotoxic stress.

### Zinc coordinately reverses the age-associated decline of the antioxidant, chaperone, proteasome, and bioenergetic network

The dominance of the ribosomal signature notwithstanding, the response to zinc was not confined to translation. As effective proteostasis requires the concerted action of redox defense, molecular chaperones, and protein degradation, we asked whether zinc reinforces these arms of the network in parallel. Our proteomics analysis confirms a broad beneficial response for proteostasis, of which example proteins are highlighted in **Figure 4A**. These include downregulation of cellular stress marker GCS-1 and upregulation of oxidative stress response proteins (SOD-1, PRDX-2), chaperones (HSP-6, PFD-2), proteins related to protein degradation (UBQL-1, PBS-3, RPT-5) as well as energy metabolism (UCR-1, ATP-4) (**Supplementary Table 1**). These zinc-responsive proteomic changes were underpinned by a coordinated transcriptional program, as analysed by RT-qPCR (**Fig. 4B-D, H-N**). Among antioxidant and redox genes, *sod-1* decreased to 0.45-fold and was restored to 1.06-fold by zinc supplementation (**Fig. 4C**; *p* < 0.001, treated versus untreated day 1), *prdx-2* was restored from 0.45- to 1.25-fold (**Fig. 4D**; *p* < 0.001), and *gcs-1*, which increased 1.54-fold in untreated aged animals (*p* < 0.01 versus day 0), was returned to 0.87-fold by zinc supplementation (*p* < 0.01 versus untreated day 1) (**Fig. 4B**). We assessed if transcriptomic and proteomic changes had a functional impact on the antioxidant capacity. Total superoxide dismutase (SOD) activity decreased significantly during aging from 17.2 to 12.5 U/mL between day 0 and untreated day 1 (**Fig. 4E**; *p* < 0.001) and zinc supplementation restored it to youthful levels (17.5 U/mL; *p* < 0.001 versus untreated day 1, not significant versus day 0). Consistent with enhanced antioxidant defense, zinc-supplemented animals were more resistant to acute oxidative challenge with paraquat, surviving significantly longer than untreated animals (**Fig. 4F**; log-rank *p* < 0.001 for 10 μM versus 0 μM, whereas 1 μM zinc conferred no significant protection). Zinc also conferred robust thermotolerance. At 35°C, 10 μM zinc could prolong survival relative to both 0 μM and 1 μM (**Fig. 4G**; log-rank *p* < 0.0001 for both comparisons), and even 1 μM afforded significant protection over non-supplemented animals. Thus, at the level of whole-animal physiology, even short-term zinc supplementation restores two hallmarks of health; antioxidant capacity and proteotoxic-stress tolerance.

**Figure 4.**
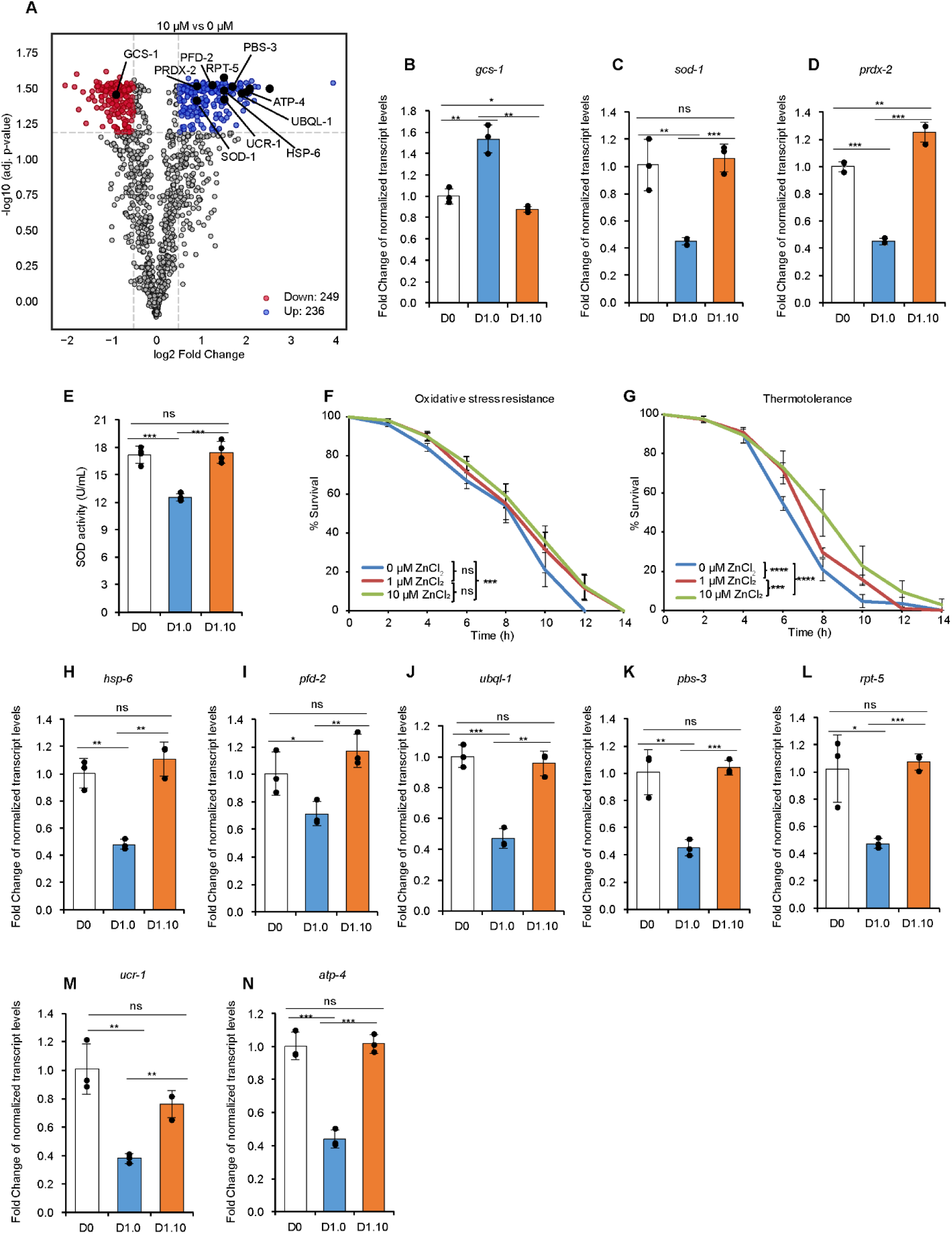
Zinc supplementation coordinately reverses the age-associated decline of antioxidant, chaperone, proteasome, and bioenergetic genes and restores stress resistance. (A) Volcano plot (10 μM versus 0 μM, day 1) with validated stress, redox, chaperone, and proteasome proteins labelled. Significantly changed proteins are colored (permutation-based FDR < 0.05 and log2 fold change ±0.5). (B-D) RT-qPCR of redox genes *gcs-1*, *sod-1*, *prdx-2*. (E) Total SOD activity (WST-1 assay) at day 0, untreated day 1 (D1.0), and zinc-treated day 1 (D1.10); activity fell with age and was restored by zinc (mean ± SD, n = 4; *p* < 0.001). (F) Oxidative-stress resistance (50 mM paraquat) of animals pre-treated 3 days with 0, 1, or 10 μM ZnCl_2_; 10 μM significantly prolonged survival (log-rank *p* < 0.001 versus 0 μM); 1 μM ns. (G) Thermotolerance at 35 °C; 10 μM ZnCl_2_ markedly prolonged survival (log-rank *p* < 0.0001 versus 0 μM and versus 1 μM). (H-N) RT-qPCR of chaperone (*hsp-6*, *pfd-2*), bioenergetic (*ucr-1*, *atp-4*), and proteasome/ubiquitin (*pbs-3*, *ubql-1*, *rpt-5*) genes. For all RT-qPCR panels, fold change is relative to day 0 baseline, normalized to *cdc-42*; mean +/- SD, *n* = 3; *\*p* < 0.05, *\*\*p* < 0.01, *\*\*\*p* < 0.001, ns - not significant.

Other analysed genes followed a similar trend where zinc-supplemented animals showed significant increases in transcript levels compared to non-supplemented animals. These included the stress-inducible mitochondrial chaperone *hsp-6* and the prefoldin subunit *pfd-2* (**Fig. 4H and I**; *p* < 0.01 and *p* < 0.01, respectively). Increases in the proteasome subunits *pbs-3* and *rpt-5* as well as *ubql-1,* which binds client proteins and shuttles them to the ubiquitination machinery, suggest an increased protein degradation capacity upon zinc supplementation (**Fig. 4J-L**; *p* < 0.001, *p* < 0.01, and *p* < 0.001, respectively). Finally, aging is characterized by a decrease in energy metabolism and our zinc-supplementation regimen was able to increase the bioenergetic genes *ucr-1* and *atp-4* (**Fig. 4M and N**; *p* < 0.01 and *p* < 0.001). The extent of this response, spanning redox defense, cytosolic and mitochondrial chaperones, the ubiquitin-proteasome system, and oxidative-phosphorylation components, indicates that zinc does not merely tune a single pathway but acts as a coordinating hub that stabilizes the proteostasis network.

### Zinc protects against polyglutamine and amyloid-beta proteotoxicity

To determine whether zinc-reinforced proteostasis translates into protection against disease-relevant protein aggregation, we turned to two established *C. elegans* models of human proteinopathy. We first examined polyglutamine (polyQ) toxicity using the AM141 strain, which expresses a 40-residue polyglutamine tract fused to YFP in body-wall muscle (Q40::YFP). This model shows progressive polyQ aggregation and motility loss during aging recapitulating the pathogenic threshold of Huntington’s disease and related polyQ disorders [23]. Late-life zinc supplementation extended the lifespan of AM141 animals, raising median survival from 5 to 7 days, and produced a comparable extension in the corresponding YFP-only control strain OW450 (from 9 to ∼11 days), indicating a benefit on both the disease and the underlying aging process (**Fig. 5A**). Quantification of muscle polyQ aggregates showed that aggregate burden increased steeply with age in untreated animals, from a baseline of ∼42 foci per animal to ∼72 by day 3, and that zinc supplementation significantly suppressed this accumulation (**Fig. 5B and C**). By day 3 of treatment, zinc-supplemented animals harbored fewer aggregates than untreated animals (∼52 versus ∼72 foci, Mann-Whitney *p* < 0.001), effectively holding aggregate load near the day 1 level. The suppression of age-driven polyQ aggregation demonstrates that zinc protects against a protein-conformational disease phenotype.

**Figure 5.**
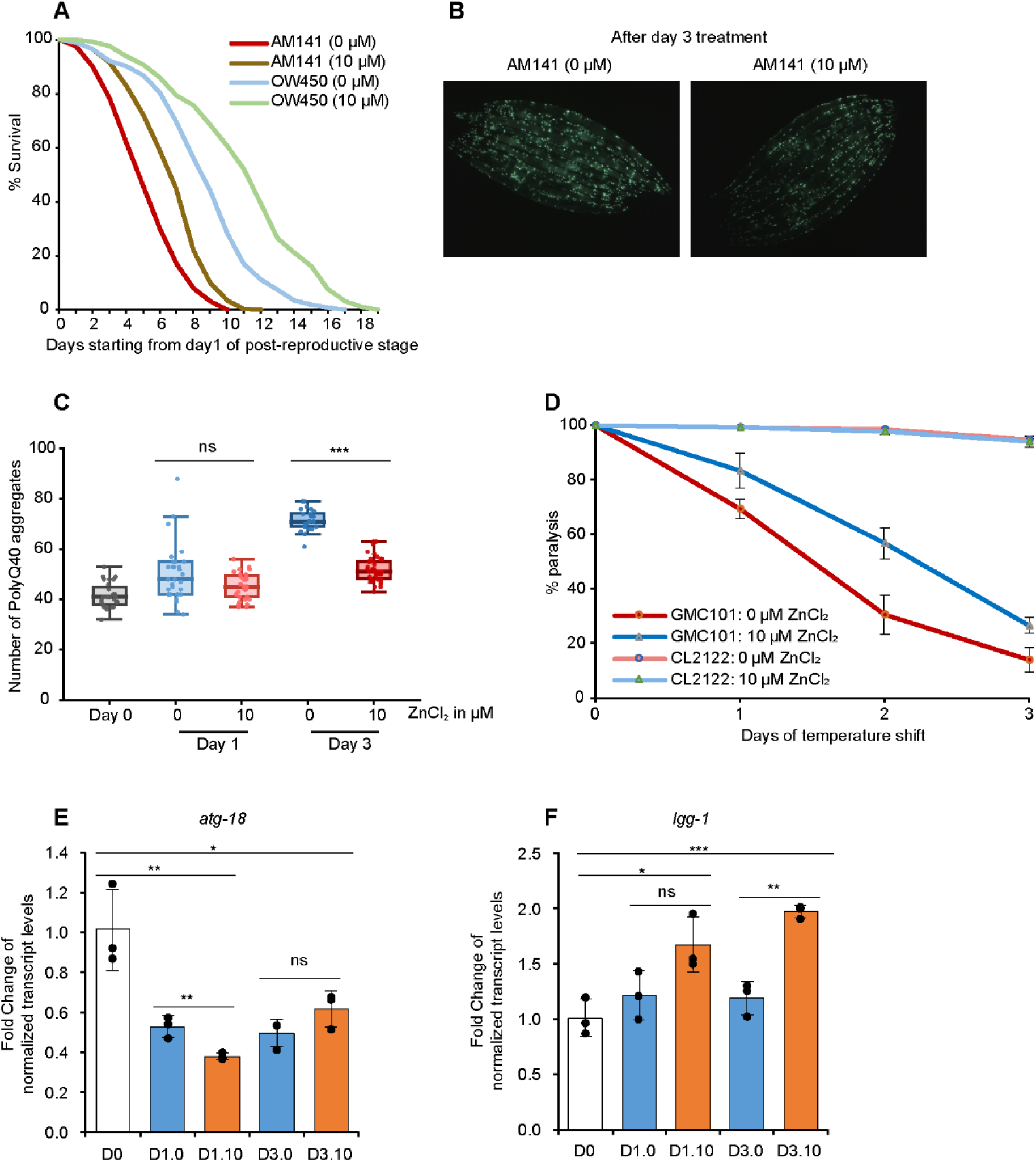
Zinc supplementation protects against polyglutamine and Aβ proteotoxicity and primes a selective autophagy response. (A) Kaplan-Meier survival of the polyQ model AM141 (Q40::YFP) and its YFP-only control OW450, each on 0 or 10 μM ZnCl_2_ from the first day of the post-reproductive stage; zinc extended median survival of both strains (AM141, 5→7 days; OW450, 9→∼11 days). (B) Representative fluorescence images of AM141 muscle polyQ aggregates after 3 days on 0 versus 10 μM ZnCl_2_. (C) Aggregate counts per animal at baseline (day 0) and after 1 or 3 days on plates supplemented with 0 or 10 μM ZnCl_2_; aggregates accumulate with age and are significantly suppressed by zinc at day 3 (Mann-Whitney with BH-FDR; *p* < 0.001; ns at day 1). (D) Cumulative paralysis of the Aβ1-42 model GMC101 and the non-Aβ control CL2122 after temperature upshift (28°C), NAMM with 0 or 10 μM ZnCl_2_; zinc significantly slowed GMC101 paralysis (Welch’s t-test, *p* < 0.05, 10 μM versus 0 μM, days 1-3), with no effect in CL2122. (E, F) RT-qPCR of the autophagy genes *atg-18* (bulk autophagosome biogenesis) and *lgg-1* (Atg8/LC3, selective cargo capture) across aging and zinc treatment; zinc induces *lgg-1* (up to ∼2.0-fold by day 3; *p* < 0.001) but does not induce *atg-18*, which is reduced by zinc at day 1 (*p* < 0.01). Mean ± SD, *n* = 3; fold change relative to day-0 baseline, normalized to *cdc-42*.

We next tested an amyloid-beta model using the GMC101 strain, which expresses human Aβ1-42 in body-wall muscle and undergoes rapid, temperature-inducible paralysis as the peptide oligomerizes and aggregates [24]. Following a temperature upshift to induce aggregation, GMC101 animals that were maintained without additional zinc were paralyzed most rapidly, whereas 10 μM zinc supplementation significantly slowed the progression of paralysis at every time point (**Fig. 5D**; two-sample Welch’s t-test, *p* < 0.05 for 10 μM versus 0 μM at days 1-3). The non-Aβ-expressing control strain CL2122 showed negligible paralysis under all conditions, confirming that the phenotype and its zinc-dependent rescue are specific to Aβ proteotoxicity. Zinc thus confers protection across two proteinopathies consistent with an action on the general proteostasis machinery.

### Zinc primes a selective autophagy response rather than bulk autophagy

Macroautophagy is a principal route for clearing aggregation-prone proteins and is required for lifespan extension by several longevity interventions [25, 26]. We asked whether zinc engages the autophagy pathway, and if so, in what mode. To this end, we measured the autophagy-related transcripts *lgg-1* (the *C. elegans* orthologue of yeast Atg8 and mammalian LC3, central to autophagosome formation and selective cargo capture) and *atg-18* (a WIPI-family factor required for bulk autophagosome biogenesis) across aging and zinc treatment (**Fig. 5E and F**). The two genes behaved oppositely. The *atg-18* gene was not induced by zinc; rather, it declined with age and was further reduced by zinc at day 1 (0.53-to 0.38-fold, *p* < 0.01 versus untreated day 1), with no difference from untreated animals at day 3 (**Fig. 5E**). In contrast, *lgg-1* was progressively induced during zinc supplementation, rising significantly above baseline at day 1 (*p* < 0.05) and reaching ∼2.0-fold by day 3 of treatment (**Fig. 5F**; *p* < 0.001 for day 3 zinc-treated versus baseline, and *p* < 0.01 versus untreated day 3). The selective induction of *lgg-1* without concomitant activation of *atg-18* indicates that zinc does not switch on indiscriminate bulk autophagy but instead primes a targeted, selective autophagy response directed toward the disposal of specific proteotoxic cargo. This mechanistic distinction may set zinc supplementation apart from interventions such as dietary restriction, which extend lifespan through broad induction of the core autophagy machinery [26]. Consequently, it provides a plausible route for zinc to facilitate the selective degradation of age-related misfolded proteins without incurring the metabolic costs of excessive bulk autophagy.

## Discussion

Age-associated declines in zinc status have been shown particularly in humans and mice models and have traditionally been studied in the context of immunosenescence, chronic inflammation and impaired antioxidant defences [27–29]. In aged mice, for example, reduced zinc status accompanies thymic involution and immune dysfunction, while dietary zinc supplementation can restore circulating zinc and ameliorate defects in thymopoiesis and inflammatory responses [30–32]. These observations have led to the prevailing view that age-associated zinc loss is largely a nutritional or systemic defect that compromises zinc-dependent physiological processes. Whether declining zinc availability also compromises core mechanisms of cellular proteostasis, a hallmark of aging, has received considerably less attention [33]. *C. elegans* is a well-established model for studying zinc homeostasis, with conserved zinc transport and sensing pathways and a defined aging trajectory [34]. We showed that whole-body metallomics revealed a pronounced decline in zinc during late post-reproductive life. This distinguishes zinc from the broader pattern of age-associated metal accumulation such as iron, calcium and magnesium previously reported in *C. elegans* [35] and confirmed by our present study. Importantly, zinc depletion occurred despite continuous dietary exposure suggesting age-dependent changes in zinc uptake, intestinal transport, intracellular buffering/storage, tissue redistribution, or zinc loss/excretion.

Our findings add an interesting perspective to the seemingly counterintuitive relationship between zinc and longevity in *C. elegans*. Previous research reported that chronic exposure to high zinc shortened lifespan and impaired healthspan, whereas zinc depletion extended lifespan and reduced protein insolubility [36]. Our results suggest that these effects may depend strongly on the animal’s initial zinc status. While excessive zinc is detrimental, low-dose supplementation was beneficial when zinc was restored after its substantial decline during aging. Thus, rather than simply being beneficial or harmful, the effects of zinc may depend on both its dose and the physiological context in which it is provided.

A substantial fraction of cellular zinc is bound to proteins, thus changes in zinc availability can have broad effects on the proteome. In yeast, zinc deficiency leads to a marked reduction in ribosomal proteins and other components of the translational machinery, together with changes in chaperones and other stress-response proteins [37]. This suggests that protein synthesis is particularly sensitive to zinc availability and that cells respond to zinc limitation, in part, by reducing their demand for zinc-containing proteins. Our findings extend this observation to aging *C. elegans*. We find that the protein synthesis machinery is a major target of age-associated zinc loss. Restoring zinc availability increases the abundance of many cytosolic ribosomal proteins, enhances protein synthesis and supports translational fidelity. Several ribosomal proteins contain structural zinc sites that contribute to their stability, making them particularly plausible targets of changes in zinc availability [17]. The ribosome may therefore represent an important zinc-containing protein pool that is vulnerable to zinc loss during aging. This possibility is further supported by recent evidence from yeast that zinc-binding cysteine residues in ribosomal proteins can undergo reversible oxidation during aging, which may promote zinc release; conversely, zinc loss may leave these cysteine residues vulnerable to oxidation [16, 17, 38].

The link between zinc, ribosomal proteins and protein synthesis raises an important question. If zinc loss affects the translational machinery, why does restoring zinc increase protein synthesis when reduced translation is generally associated with longevity? Indeed, lowering translation through TOR inhibition, reduced initiation or depletion of ribosomal proteins can extend lifespan in *C. elegans* and other organisms [7, 39]. More recently, reducing ribosome biogenesis itself has also been shown to promote longevity [40]. Here, restricting Pol I activity and pre-rRNA synthesis extends lifespan in *C. elegans*, whereas increased rRNA synthesis accelerates metabolic decline and aging. Our findings suggest that this apparent contradiction may reflect an important difference between how much protein is made and how well it is made. Zinc supplementation increased ribosomal protein abundance and protein synthesis without increasing pre-rRNA or mature rRNA levels, rather we observed a decrease in the latter. Thus, zinc does not appear to promote protein synthesis by simply producing more ribosomes. Instead, it may improve the functional capacity of existing ribosomes. This interpretation is consistent with evidence that translation fidelity is an important determinant of healthy aging, independent of overall translation rate [13, 41]. Increasing translation accuracy through a single RPS23 mutation extends lifespan and improves stress resistance in yeast, *C. elegans* and flies without increasing protein synthesis, while comparative studies in mammals have linked higher translation fidelity with longer lifespan [9]. Thus, although reducing overall translation can promote longevity, restoring the structural integrity of ribosomes may have a similar outcome. Zinc may therefore improve productive protein synthesis while limiting the generation of aberrant proteins, potentially reducing the burden placed on downstream protein quality-control pathways [42–44]. The protection we observe against both polyglutamine and Aβ proteotoxicity is consistent with such a broader effect on proteostasis. Zinc appears to improve the cellular environment in which different forms of proteotoxic stress arise and are managed. The accompanying increase in *lgg-1* expression further suggests that zinc may influence protein clearance, although increased *lgg-1* alone does not establish increased autophagic flux [45]. Whether this response is a consequence of improved translational proteostasis or represents an independent arm of the zinc response awaits future testing. Together with the changes in protein abundance of chaperones and the proteasome that we observed, however, the data point toward a coordinated effect of zinc on several components of the proteostasis network rather than activation of a single stress pathway.

Together, our findings suggest that age-related zinc loss may weaken proteostasis by affecting the quality of protein synthesis and, consequently, the wider proteostasis network. Zinc repletion may therefore help maintain protein homeostasis by supporting the functional capacity of translation. Maintaining adequate zinc availability in late life can thus be an important component of proteostatic resilience.

## Materials and methods

### *C. elegans* strains and maintenance

All *C. elegans* strains were obtained from the *Caenorhabditis* Genetics Center (CGC), funded by the NIH Office of Research Infrastructure Programs (P40 OD010440). Wild-type *C. elegans* (Bristol N2) was used for all lifespan, healthspan, biochemical, and proteomic experiments. For polyglutamine aggregation quantification and lifespan assays, strain AM141 [rmIs133(unc-54p::Q40::YFP)], expressing a 40-residue polyglutamine tract fused to YFP in body-wall muscle, was used for aggregation assays, and OW450 [rmIs126(unc-54p::YFP) V], expressing YFP alone in body-wall muscle, was used as the corresponding non-aggregating control strain. For the amyloid-beta paralysis assay, GMC101 [dvIs100(unc-54p::Abeta1-42::unc-54 3’-UTR + mtl-2p::GFP)] was used as the paralysis model, with CL2122 [dvIs15(unc-54(vector) + mtl-2p::GFP)] as a non-Abeta-expressing transgenic control strain. Worms were maintained at 20 °C on nematode growth medium (NGM) seeded with *Escherichia coli* strain HB101 according to standard procedures [46, 47].

### Zinc supplementation, NAMM medium, and chelator treatment

Age-synchronised populations were obtained by hypochlorite (alkaline bleach) lysis of gravid adults. Synchronised eggs were plated on standard NGM plates seeded with *E. coli* strain HB101 and cultured until post-reproductive day 6 at 20 °C. Adults were transferred to fresh plates daily during the reproductive period to separate them from progeny well before the initiation of any treatment. At post-reproductive day 6, worms were transferred to Noble Agar Minimal Medium (NAMM; 1.7% Noble agar, 5 μg/mL cholesterol) supplemented with ZnCl_2_ at final concentrations of 0 μM (vehicle control), 1 μM, 10 μM, 100 μM, 1 mM, and 10 mM. Cholesterol was added alongside the ZnCl_2_ to molten NAMM below 55 °C immediately before pouring. 10 μM ZnCl_2_ was identified as optimal and used for all subsequent mechanistic experiments. Every NAMM plate was used within 2 weeks of pouring. For zinc chelation experiments, the membrane-permeant chelator TPEN [N,N,N’,N’-tetrakis(2-pyridylmethyl)ethylenediamine] was included at 200 μM final concentration, alone or with 10 μM ZnCl_2_.

### Lifespan analysis

Lifespan assays were performed by transferring age-synchronized post-reproductive day 6 wild-type (N2) worms to NAMM plates supplemented without or with different concentrations of ZnCl_2_ (1 μM, 10 μM, 100 μM, 1 mM, and 10 mM). At least 100 worms were scored per condition, across three independent biological replicates. For AM141 and OW450, worms were cultured under standard conditions until day 7 of adulthood, at which point they were transferred to plates with or without 10 μM ZnCl_2_. Worms were scored every day and considered dead when unresponsive to gentle prodding. Animals showing internal hatching, extruded gonads, or that crawled off the plate were censored. Survival was analysed by the Kaplan-Meier method and compared by the log-rank (Mantel-Cox) test. To test whether the zinc benefit depends on bacterial metabolic activity rather than acting directly on the host, we performed lifespan assays using heat-killed *E. coli* HB101. Bacteria grown to 10^9^ CFU/mL were heat-killed at 70°C for 2 h, and inactivation was confirmed by the absence of growth on LB agar plates.

### Locomotion analysis

#### Using WMicrotracker

Neuromuscular function was assessed by quantifying locomotory activity using the WMicrotracker ONE system (PhylumTech, Santa Fe, Argentina), which detects worm movement through transient interruptions of infrared microbeams, recorded as phototransistor output. Worms were cultured until post-reproductive day 6, after which plates were washed with 20 mM 3-(N-morpholino)propanesulfonic acid (MOPS) buffer (pH 7.0) and worms were washed multiple times with MOPS buffer through gravity settlement to remove residual bacteria. Worm populations were adjusted to a density of 10 worms/10 μL in MOPS buffer, and 50 μL volumes were dispensed into individual wells of flat-bottom 96-well microtiter plates (Costar). A concentration-dependent locomotion assay was performed using ZnCl_2_. Fifty microlitres of ZnCl_2_ solution were added to each well to yield final concentrations of 0, 1 μM, 10 μM, 100 μM, 1 mM, and 10 mM in a final well volume of 100 μL. For zinc chelation experiments, TPEN (200 μM final concentration) was co-administered with ZnCl_2_ across the range of concentrations. Locomotory activity was monitored continuously over 72 h, with all conditions normalized to their respective baseline value to control for variability.

#### Using WormLab tracking

Synchronized worm populations (∼20 worms) were transferred, without food, onto 35 mm dishes containing clear agarose medium prepared as standard NGM but with agar replaced by agarose as the solidifying agent (3 g/L NaCl, 2.5 g/L peptone, 2% w/v agarose), to permit optical clarity for video capture. Worm movement was recorded for 10 min at post-reproductive day 6 (designated day 0), after which worms were maintained without or with 10 μM ZnCl_2_ and re-recorded at day 1 (24 h) and day 3 (72 h) of treatment. Videos were analyzed using WormLab software (v2023). Images were captured in dark-worms-on-light-background mode using a flexible, video-specific threshold. Gradient correction, smoothing, illumination modelling, and hole-filling were disabled. The following settings were used for automated worm tracking, Threshold Level: Flexible, based on the video; Image Mode: Dark Worms on a Light Background; Gradient Correction (Off); Smoothing (Gaussian Filtering) (Off); Use Illumination Model (Off); Fill Holes (Off) Detection Parameters: Detect worms at the edge of the image: (On); Area: [20,10000]; Length: [50,10000]; Width: [0,10000]; Width/Length Ratio: [0.05,0.25]; Detection Frequency: 90 (to prevent too many tracks to be generated); Use Length Fitting: (Off); Use Width Fitting: (On); Use Whole Plate Mode: (Off); Min Contour Fit: 0.75; Width Regularity Threshold: 0.8; Spline Base Size: 8; Spinal Axis Sample: 59; Width Fit Frequency: 1; Fitting Iterations: 80;Tracking Parameters: Start Frame: 0; End Frame: depends on the video length; Use Back Tracking: (On); Track worms at the edge of the image: (On); Max Tracked Hypotheses: 5; Frames worms can touch boundary: 16; Frames worms can overlap: 267; Position Tolerance: 0.3; Shape Tolerance: 0.25. Minimal track-length filtering and Kaplan smoothing were applied throughout. Average speed and mobility metrics were exported for behavioural analysis. Only tracks exceeding 700 frames were retained for downstream analysis, and tracks were averaged frame-wise within each biological replicate. Three biological replicates with 50-60 individual worms were observed. Each biological replicate was analyzed by averaging the filtered tracks at every frame, for the whole recorded duration. All the biological replicates were averaged frame wise to generate the histogram. For statistical analysis, biological replicates were individually analysed to decompose the speed values into mode from histogram, positive and negative metrics. Levene’s test was used to assess variance homogeneity, followed by Student’s t-test assuming equal variances for biologically relevant pairwise condition comparisons.

### Superoxide dismutase (SOD) activity assay

SOD activity was measured using a WST-1-based colorimetric assay, adapted from [48]. Briefly, 50 mM phosphate buffer (pH 7.8) was prepared from K_2_HPO_4_ (0.0428 M) and KH_2_PO_4_ (0.0072 M). Assay buffer was made by combining phosphate buffer with hypoxanthine (10 mM stock). Reaction Mix A consisted of assay buffer, WST-1 (10 mM), and catalase (2 mg/mL). Reaction Mix B consisted of freshly diluted xanthine oxidase. Standards (bovine Cu/Zn-SOD) and samples (20 μL) were loaded into a clear, flat-bottom 96-well plate. Reaction Mix B was added to Reaction Mix A immediately before use, and 180 μL of the combined mixture was rapidly dispensed into each well. Plates were incubated at 37 °C, and absorbance was read at 438 nm after 30 min using a BioTek Synergy H1 microplate reader (Agilent). SOD activity was calculated by interpolation against the standard curve.

### Ribosomal RNA analysis

Post-reproductive day 6 adult *C. elegans* (day 0 of treatment) were maintained on NGM (day 0 baseline) or transferred to NAMM medium with 0 μM or 10 μM ZnCl_2_, and collected at day 1 and day 3 (approximately 1000 worms/replicate, three biological replicates/condition). Worm pellets were resuspended in 1 mL TRIzol (Invitrogen) and lysed by five freeze-thaw cycles (liquid nitrogen/37°C) followed by five vortex cycles (approximately 3200 rpm, 30 s on/off). RNA was extracted with chloroform (200 μL/mL TRIzol; 12,000 *g*, 15 min, 4°C, aqueous phase collected twice), precipitated with isopropanol (500 μL/mL TRIzol, -20°C overnight; 12,000 *g*, 10 min, 4°C), washed twice with 70% ethanol (7,500 *g*, 5 min, 4°C), and resuspended in 20 μL DEPC water. RNA concentration was measured and equalized across samples. Samples were DNase-treated (RQ1 RNase-Free DNase, Promega; 1 U/μg RNA, 37°C, 10 min; inactivated with DNase Stop Solution, 65°C, 10 min). rRNA quality and quantity were assessed using an Agilent Femto Pulse System (Ultra Sensitivity RNA Analysis Kit).

### SUnSET (surface sensing of translation) assay

Age-synchronized, post-reproductive day 6 worms were cultured and transferred to NAMM plates containing 0 μM or 10 μM ZnCl_2_, and collected on day 1 and day 3 of treatment. Worms were homogenized in lysis buffer (10 mM Tris-HCl, pH 7.5, 40 mM NaCl, 30 mM MgCl_2_, 1 mM PMSF, and protease inhibitors [1 μM pepstatin, 10 μg/mL aprotinin, 10 μM leupeptin]) containing puromycin (from a 10 mg/mL stock, final concentration 4 mM) using a Mixer Mill (MM 400, Retsch GmbH, Haan, Germany) for 15 min at 30 Hz. Lysates were cleared by centrifugation, and protein concentrations were determined using the bicinchoninic acid (BCA) assay. Equal amounts of protein were loaded onto a 12.5% SDS-PAGE gel and transferred to a PVDF membrane for Western blotting. Total protein was visualized by Coomassie staining of the membrane. For puromycin detection, membranes were blocked in 5% (w/v) milk in 1xTBS and incubated overnight at 4°C with mouse anti-puromycin primary antibody (clone PUR333.10, EPRO Science) at 1:20,000 dilution. After washing, blots were incubated with anti-mouse IgG secondary antibody (1:10,000 dilution) for 2 h. Following three washes with 1xTBS, chemiluminescent detection was performed using ECL. Band intensities for puromycin and total Coomassie-stained protein were quantified using ImageJ, and puromycin incorporation was expressed as the puromycin/Coomassie signal ratio for each condition. Data are presented as mean +/- SD from *n* = 3 independent experiments.

### Bright-field imaging of body posture

Post-reproductive day 6 worms maintained on NGM plates were transferred to HB101-seeded NAMM plates without supplementation (0 μM ZnCl_2_) or with 10 μM ZnCl_2_ supplementation and cultured until day 3 of treatment. At this endpoint, individual worms from each condition were picked onto NAMM plates for assessment of sinusoidal or uncoordinated movement on their respective ZnCl_2_ concentrations and imaged using a Zeiss stereomicroscope (Jena, Germany).

### Reverse transcription real-time PCR (RT-qPCR)

Total RNA was extracted (see the “Ribosomal RNA” analysis section) and subjected to DNase I treatment. Reverse transcription was performed using an equal concentration of RNA (1 μg) for each condition and its replicate according to the manufacturer’s instructions (Cat. no. 205313, Qiagen). Following reverse transcription, cDNA was diluted 5-fold in nuclease-free water prior to use. The reactions for real-time PCR were prepared in a 20 μL final volume containing 1 μL diluted cDNA, 1 μL forward primer and 1 μL reverse primer (each from a 10 μM stock, yielding a final concentration of 0.5 μM each; Supplementary Table 3), 10 μL RT-PCR Mix SYBR C (Cat. no. 2008-100C; A&A Biotechnology), and 7 μL nuclease-free water. Samples were loaded onto a 96-well plate along with negative (no-template) controls. Real-time PCR was performed using a LightCycler 480 System (Roche, Basel, Switzerland). The reaction program included an initial denaturation step for 5 min (95 °C) followed by 40 cycles of amplification (95 °C for 30 s, 57 °C for 20 s, and 72 °C for 20 s), followed by melt-curve analysis. Primers used for the analysis are listed in Supplementary Table 3. Relative expression of the pre-rRNA precursor was normalized to *snb-1*, while relative expression of other genes was normalized to *cdc-42*, chosen to align with the reference gene used in the corresponding proteomics dataset. Relative expression was calculated using the comparative 2^−ΔΔCt^ method. For each replicate, ΔCt was calculated as ΔCt = Ct(target) − Ct(reference gene). ΔΔCt was then calculated relative to the mean ΔCt of the appropriate control condition: for comparisons against the untreated baseline, ΔΔCt = ΔCt(sample) − mean ΔCt(baseline condition); for treatment-versus-vehicle comparisons at matched timepoints, ΔΔCt = ΔCt(treated) − mean ΔCt(untreated control, same timepoint). Fold change was calculated as 2^−ΔΔCt^ and reported as mean +/- SD from three biological replicates. Statistical significance between conditions was assessed using an unpaired, two-tailed Student’s *t*-test; *p* < 0.05 was considered statistically significant.

### Proteomic sample preparation

Age-synchronized, post-reproductive day 6 animals (considered as day 0) treated with 0 or 10 μM ZnCl_2_ were harvested at day 1 and day 3 post-treatment (four biological replicates per condition). Worms were washed three times with M9 buffer to remove residual bacteria and snap-frozen. Samples were prepared according to the protocol of [49]. After thawing, samples were washed twice with 100 mM Tris buffer (pH 8.5), resuspended in 200 μL of 8 M urea lysis buffer (8 M urea, 100 mM Tris, pH 8.5), and lysed by probe sonication (4-6 pulses of 10 s each). Lysates were clarified by centrifugation (12,000 rpm, 15 min, 4 °C). Protein concentrations of supernatants were determined by Bradford assay (ROTIQuant, Cat. no. K015.3, Carl Roth GmbH) with bovine serum albumin as a calibrating protein. For each sample, 50 μg of protein was diluted 8-fold with 100 mM Tris-HCl (pH 8.5) and digested by overnight incubation (16 h, 37 °C, 400 rpm) with sequencing-grade modified trypsin (1:50 enzyme-to-protein ratio, w/w) in the presence of 10 mM tris(2-carboxyethyl)phosphine (TCEP) and 40 mM chloroacetamide (CAA). Digestion was quenched by acidification with trifluoroacetic acid (TFA) to a final concentration of 0.5-2% (pH approximately 2). Samples were snap-frozen on dry ice and stored at -80°C prior to tandem mass tag (TMT) labelling.

### TMT labelling and fractionation

The resulting peptides were labelled using an on-column TMT labelling protocol. Stage tips were packed with three punches of C18 mesh (Affinisep) with a 16-gauge blunt-end needle. Resin was conditioned with 150 μL methanol, followed by 100 μL 50% acetonitrile (ACN)/0.1% formic acid (FA), and equilibrated twice with 150 μL 0.1% FA. The digested peptides (10 μg from each sample) were loaded by spinning at 1200 *g* until the entire digest had passed through. The bound peptides were washed twice with 150 μL 0.1% FA. 200 μL of TMT reagent in 50 mM phosphate buffer pH 8 (approximately 80 μg of TMT reagent resuspended in 2 μL 100% ACN and diluted with 200 μL of 50 mM phosphate buffer pH 8) was loaded over the C18 resin at 300 *g* until the entire solution had passed through. The phosphate buffer and residual TMT were washed away three times with 150 μL 0.1% FA. Peptides were eluted with 60 μL 60% ACN/0.1% FA. TMT-labelled samples were combined into a single TMT sample and concentrated using a SpeedVac concentrator.

Peptides in the combined sample were separated into 8 fractions (fractions 1, 2, and 3 were combined) by offline basic reversed-phase chromatography using the Pierce High pH Reversed-Phase Peptide Fractionation Kit (Thermo Fisher Scientific).

### Liquid chromatography-mass spectrometry analysis

Prior to liquid chromatography-mass spectrometry (LC-MS/MS) measurement, the peptide fractions were resuspended in 0.1% TFA and 2% acetonitrile in water. Chromatographic separation was performed on a Dionex UltiMate 3000 nano-LC system using an Easy-Spray Acclaim PepMap column (50 cm length x 75 μm inner diameter; Thermo Fisher Scientific) at 55 °C by applying 120 min acetonitrile gradients in 0.1% aqueous formic acid at a flow rate of 300 nL/min.

The Q Exactive HF-X mass spectrometer was operated in TMT mode with survey scans acquired at a resolution of 60,000 at m/z 200. Up to 15 of the most abundant isotope patterns with charges 2-5 from the survey scan were selected with an isolation window of 0.7 m/z and fragmented by higher-energy collisional dissociation with a normalized collision energy of 32, while the dynamic exclusion was set to 30 s. The maximum ion injection times for the survey scan and MS/MS scans (acquired at a resolution of 45,000 at m/z 200) were 50 and 150 ms, respectively. The ion target value for MS was set to 3e6 and for MS/MS to 1e5, and the minimum AGC target was set to 1e3.

### Proteomics data processing

LC-MS/MS spectra were searched against the *C. elegans* reference proteome (UniProt Proteome ID UP000001940) using MaxQuant (v2.2.0.0). Normalization using weighted ratio to reference channel 1 (the mixture of all samples) was applied. Carbamidomethylation of cysteine was set as a fixed modification, and methionine oxidation and protein N-terminal acetylation as variable modifications. Digestion specificity was set to Trypsin/P. Match-between-runs was enabled. The processed data (proteinGroups.txt, reporter intensities) were imported into Perseus (v2.0.6.0) for downstream analysis. Protein groups annotated as reverse hits, identified only by site, or potential contaminants were removed, and only proteins identified by at least one unique peptide were retained. Reporter intensities were log2-transformed. Proteins were required to have valid values in at least 75% of samples in each group; missing values were imputed from a normal distribution. Data were normalized by median subtraction across samples. Differentially abundant proteins were identified by two-sided Student’s *t*-test with permutation-based FDR correction (FDR = 0.05).

### Bioinformatic analysis of proteomics data

Annotations for detected proteins were obtained by searching the *C. elegans* reference proteome (UniProt Proteome ID UP000001940, downloaded 06.06.2025). Annotations for the zinc-binding proteome were obtained by searching for "zinc" or "ZN" within Gene Ontology Molecular Function (GOMF), Cofactor, Binding Site, Zinc finger, and Keywords. Annotations for ribosomal proteins were obtained by searching for "ribosomal protein" in Protein names. Annotations were manually curated. Proteins with CXXC motifs were identified by custom code in Python 3.10 based on protein sequences downloaded from UniProt (UP000001940, downloaded 06.06.2025). Visualization was performed using the matplotlib and seaborn libraries in Python 3.10.

Functional enrichment analysis was performed using STRING (string-db.org; accessed 26.08.2025) with the whole *C. elegans* proteome as background and significantly altered proteins defined by log2 fold change below -0.5 or above 0.5 for down- and up-regulated sets, respectively (FDR <= 0.05). Overrepresented terms were searched across GOMF and were visualized using matplotlib and seaborn in Python 3.10. STRING protein-protein interaction networks were analyzed with Cytoscape 3.7.0 using the STRING plugin, and MCL clustering was performed with an inflation parameter of 3. Volcano plots and rank plots were generated with Python 3.10 using pandas, matplotlib and numpy.

### Stress resistance assays

Post-reproductive day 6 adult worms were pre-treated on NAMM plates supplemented with 0, 1, or 10 μM ZnCl_2_ at 20 °C for 3 days. For the thermotolerance assay, pre-treated worms were transferred onto fresh NAMM plates containing the corresponding zinc concentration and shifted to 35°C. Survival was scored every 2 h until all worms in each condition were dead. For the oxidative stress resistance assay, pre-treated worms were individually transferred onto NAMM plates supplemented with 50 mM paraquat (methyl viologen dichloride hydrate, 98%; Cat. no. 856177; Merck, Darmstadt, Germany) and the corresponding zinc concentration, and survival was similarly scored every 2 h until all worms were dead. For both assays, each condition comprised 100 worms per biological replicate across three biological replicates.

### Measurement of zinc content by ICP-OES

For the zinc supplementation experiment, post-reproductive day 6 worms pre-cultured on NGM plates were transferred to NAMM plates supplemented with 0 or 10 μM ZnCl_2_ and collected at day 1, day 3, and day 5 of treatment. For the aging time course, worms were cultured on standard NGM plates and collected at day 1 of adulthood through day 16 at intervals of day 1, 4, 7, 10, 13, and 16 (no zinc supplementation). For both experiments, worms were washed 5 times in Milli-Q water to remove residual bacteria and surface-bound zinc, then pelleted (approximately 500 worms per biological replicate) and frozen in liquid nitrogen. For ICP-OES analysis, samples were thawed and digested in ultrapure nitric acid (Cat. no. 1.00441, Merck) at 80°C for 2 h. Digested samples were diluted with 1% (v/v) nitric acid prior to metal measurement. Elemental analysis was performed on an iCAP Pro ICP-OES instrument (Thermo Fisher Scientific) in aqueous-axial-iFR mode, equipped with an iSC-65 autosampler and a Torch Duo (Slot) Rev 02 argon plasma torch, set at 1,250 W RF power, using Qtegra software. An external calibration curve was recorded with an ICP single-element standards mix in 1% (v/v) nitric acid, and the sample was introduced via a peristaltic pump. The wavelengths for sulfur (182.034 nm), zinc (206.200 nm), manganese (257.610 nm), iron (259.940 nm), calcium (184.006 nm), copper (224.700 nm) and magnesium (285.213 nm) were selected to minimise overlapping emissions. Results were calculated from ppb data, normalized to sulfur content and expressed as the “metal”/ S ratio (μM “metal”/ μM S).

### Polyglutamine aggregation quantification

AM141 and OW450 worms were aged to day 7 of adulthood (first day of the post-reproductive stage) and transferred to NAMM plates with or without 10 μM ZnCl_2_. Worms were imaged by fluorescence microscopy at day 0 (baseline), day 1, and day 3 post-treatment to visualize polyQ aggregates in body-wall muscle, across three independent biological replicates. Aggregate quantification was performed using a custom Fiji/ImageJ macro that batch-processed TIFF images across each experimental folder images. Images were converted to RGB and thresholded using the Color Threshold tool (RGB color space; red channel 70–255, green and blue channels unrestricted at 0–255); the resulting selection was converted to a binary mask, and aggregates were identified and counted using the Analyze Particles function (size: 0-infinity; circularity: 0.50-1.00). Aggregate counts were compiled and exported for downstream statistical analysis. Data are presented as individual data points with mean +/- SD.

### Amyloid-beta (Abeta) paralysis assay

GMC101 [dvIs100(unc-54p::Abeta1-42::unc-54 3’-UTR+ mtl-2p::GFP)] animals, which express human Aβ1–42 in body-wall muscle, and CL2122 [dvIs15(unc-54(vector) + mtl-2p::GFP)], a non-Aβ -expressing transgenic control strain, were synchronized and cultured in parallel on NGM plates at 20 °C. At post-reproductive day 6, worms were transferred to NAMM plates supplemented with 0 μM or 10 μM ZnCl_2_ and maintained at 20 °C for 48 h. Aβ aggregation was then induced by shifting all plates to 28 °C, with worms remaining throughout on their respective plates. Paralysis was scored at 24, 48, and 72 h after the temperature shift by gently prodding animals with a platinum pick; at each interval, previously-moving worms were re-scored and classified as still moving, newly paralyzed (failed to move the body away from the stimulus despite retaining head movement), or censored. Cumulative paralysis rate at each timepoint was calculated as the cumulative number of paralyzed worms divided by the sum of cumulative paralyzed and still-moving worms, with censored worms excluded. Each of the four groups (two conditions x two strains) comprised approximately 100 worms per replicate across three independent biological replicates. Statistical comparisons between conditions were performed on cumulative paralysis rates using an unpaired, two-tailed Welch’s *t*-test (unequal variance). Data are presented as mean +/- SD.

### Paromomycin-induced proteotoxic stress and survival assay

At post-reproductive day 6, wild-type (N2) worms were transferred to NAMM plates supplemented with TPEN (200 μM) alone, paromomycin (0.5 mM) alone, paromomycin (0.5 mM) + ZnCl_2_ (10 μM), or paromomycin (0.5 mM) + ZnCl_2_ (10 μM) + TPEN (200 μM), yielding four treatment groups: TPEN alone, paromomycin only, Zn + paromomycin, and TPEN + Zn + paromomycin. At the post-reproductive day 5 stage, the worms in the two zinc-containing groups received a 24 h pre-treatment with 10 μM ZnCl2 prior to their transfer to experimental plates, while worms in the TPEN-alone and paromomycin-only groups received no zinc pre-treatment. Each group comprised approximately 50 worms per replicate across three independent biological replicates, each performed in three technical replicates. Proteotoxic stress was induced by heat-shocking all plates at 35°C for 3 h, with worms remaining on their respective plates. Immediately after heat shock, plates were returned to 20 °C and allowed to equilibrate for 4 h before the first scoring time point. Survival was scored at each interval by gently prodding animals with a platinum pick; previously-alive worms were re-scored as still alive, newly dead (failed to respond to stimulus), or censored (lost from the plate due to crawling off the agar or desiccation). Cumulative percent survival at each time point was calculated as the number of worms remaining alive divided by the initial population size, adjusted for censored animals, and normalized to the t0 baseline. Data are presented as mean +/- SD at each time point.

### Statistical analysis

All experiments were performed in at least three independent biological replicates unless stated otherwise, and data are presented as mean +/- standard deviation (SD). Analyses were performed in GraphPad Prism and in Python 3.10 (SciPy, statsmodels, scikit-posthocs), and *p* < 0.05 was considered statistically significant. Significance is denoted throughout as follows: one asterisk, *p*< 0.05; two asterisks, *p* < 0.01; three asterisks, *p* < 0.001; four asterisks, *p* < 0.0001; and ns, not significant.

Survival and paralysis data (Kaplan-Meier lifespan curves, paraquat oxidative-stress and thermotolerance assays, and the paromomycin proteotoxic-stress survival assay) were analysed by the log-rank (Mantel-Cox) test. For assays with three biological replicates, replicates were treated as independent strata and combined by the Mantel-Haenszel method to account for replicate-to-replicate variation; the corresponding pooled test (all animals combined into a single curve per condition) gave concordant results. Aβ paralysis (cumulative % paralysis per biological replicate) was compared between 0 and 10 μM ZnCl2 at each day by unpaired, two-tailed Welch’s t-test (n = 3).

For the ZnCl_2_ dose-response and ZnCl_2_ + TPEN locomotion assays, and for the AM141 polyQ aggregate-count data, normality (Shapiro-Wilk) and homogeneity of variance (Levene’s test) were first assessed per group. As one or more groups deviated from normality and/or exhibited unequal variances in these datasets, non-parametric tests were used: a Kruskal-Wallis omnibus test across all conditions, followed by Dunn’s post-hoc test (locomotion) or targeted pairwise Mann-Whitney U tests (aggregate counts) for comparisons of interest, in each case with Benjamini-Hochberg false-discovery-rate (FDR) correction. Kruskal-Wallis, Dunn’s, and Mann-Whitney U tests were computed in Python (SciPy/scikit-posthocs).

Quantitative real-time PCR fold changes, SOD activity, ICP-OES metal-to-sulfur ratios, and SUnSET puromycin/Coomassie ratios were compared using unpaired, two-tailed Student’s t-tests (equal variances assumed). The zinc supplementation ICP-OES time course and the metal aging time course were each tested against the relevant control (matched untreated condition, or day 1 baseline) by unpaired two-tailed *t*-test (*n* = 3 biological replicates per group).

For the proteomics dataset, differentially abundant proteins were identified by two-sided Student’s t-test with permutation-based FDR correction (FDR = 0.05) in Perseus. Functional-enrichment significance (GO, KEGG, Reactome, STRING) was assessed by the STRING platform’s hypergeometric test with Benjamini-Hochberg FDR correction (FDR <= 0.05).

## DATA AVAILABILITY

The mass spectrometry proteomics data have been deposited to the ProteomeXchange Consortium via the PRIDE [50] partner repository with the dataset identifier PXD084538 and 10.6019/PXD084538. Other data underlying this article are available in the article and in its supplementary material.

## ACKNOWLEDGMENTS

Proteomic measurements were performed at the Proteomics Core Facility, IMol Polish Academy of Sciences. Analysis of total RNA using Agilent Femto Pulse System was performed in the DNA Sequencing and Synthesis Facility, Institute of Biochemistry and Biophysics, PAS. Some strains were provided by the CGC, which is funded by NIH Office of Research Infrastructure Programs (P40 OD010440).

## AUTHOR CONTRIBUTIONS

AK – data curation, investigation, formal analysis, methodology, validation, visualization, writing – original draft; KPL – data curation, formal analysis, methodology, writing – review and editing; KJ – data curation, formal analysis, visualization, writing – review and editing; RM, MM – investigation; SG – investigation, visualisation, writing – review and editing; KJW – supervision, writing – review and editing; UT – conceptualization, supervision, funding acquisition, writing – original draft. All authors approved the final version of the manuscript.

