## Supplementary files for "Zinc preserves translational proteostasis to counteract aging and proteotoxicity in *Caenorhabditis elegans*"

### CONTENT

**Supplementary Figure 1 (Related to Figure 1).** Zinc supplementation extends lifespan on metabolically inactive bacteria and improves fine-scale locomotion, in a zinc-dependent manner.

**Supplementary Figure 2 (Related to figure 2**). Rank plots and STRING protein-protein interaction network analysis of the zinc-responsive proteome.

**Supplementary Table 1.** Proteomic analyses performed from total worm extracts from N2 (wild-type) worms that were grown to day 6 of adulthood under standard culturing conditions. The worms were then supplemented with or without 10 µM $ZnCl_2$ for 24 or 72 hours.

**Supplementary Table 2.** GO enrichment analysis.

**Supplementary Table 3.** RT-qPCR primers used in this study.

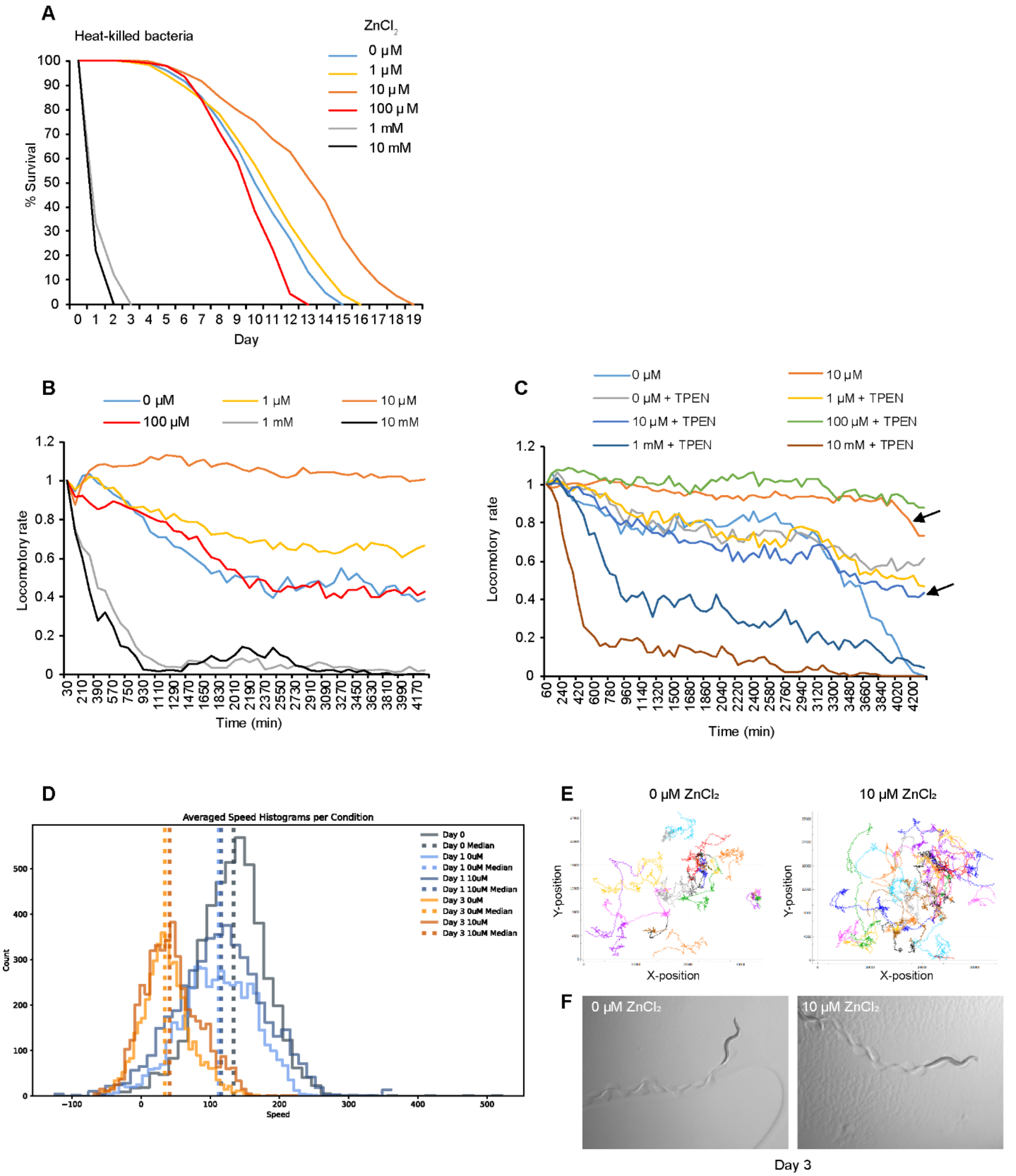


**Supplementary Figure 1. Zinc supplementation extends lifespan on metabolically inactive bacteria and improves fine-scale locomotion, in a zinc-dependent manner.** (A) Kaplan-Meier survival of N2 animals on heat-killed *E. coli* HB101 across the $ZnCl_2$ dose range; 10 µM extended median survival (11 > 14 days) whereas $\geq$ 1 mM was lethal (log-rank versus 0 µM + HK: 1 µM $p < 0.05$; 10, 100 µM and 1, 10 mM $p < 0.0001$). As the bacteria are metabolically inactive, the benefit reflects an action on the host rather than on bacterial zinc metabolism. (B) Raw single-worm locomotory-rate time courses over 72 h across the $ZnCl_2$ dose range and (C) with co-administered TPEN (WMicrotracker); arrows in (C) mark the divergence of the 10 µM and TPEN-containing conditions. (D) Averaged instantaneous-speed histograms per condition (WormLab solid-substrate tracking) at day 0, and at day 1 and 3 on 0 or 10 μM $ZnCl_2$, with per-condition medians (dashed lines). (E) Mid-position data for all tracks without filters to highlight crawling behaviour across time and conditions. Representative tracks of three biological replicates after 3 days of supplementation. (F) Representative bright-field images of body posture at day 3 of treatment on 0 versus 10 μM $ZnCl_2$.

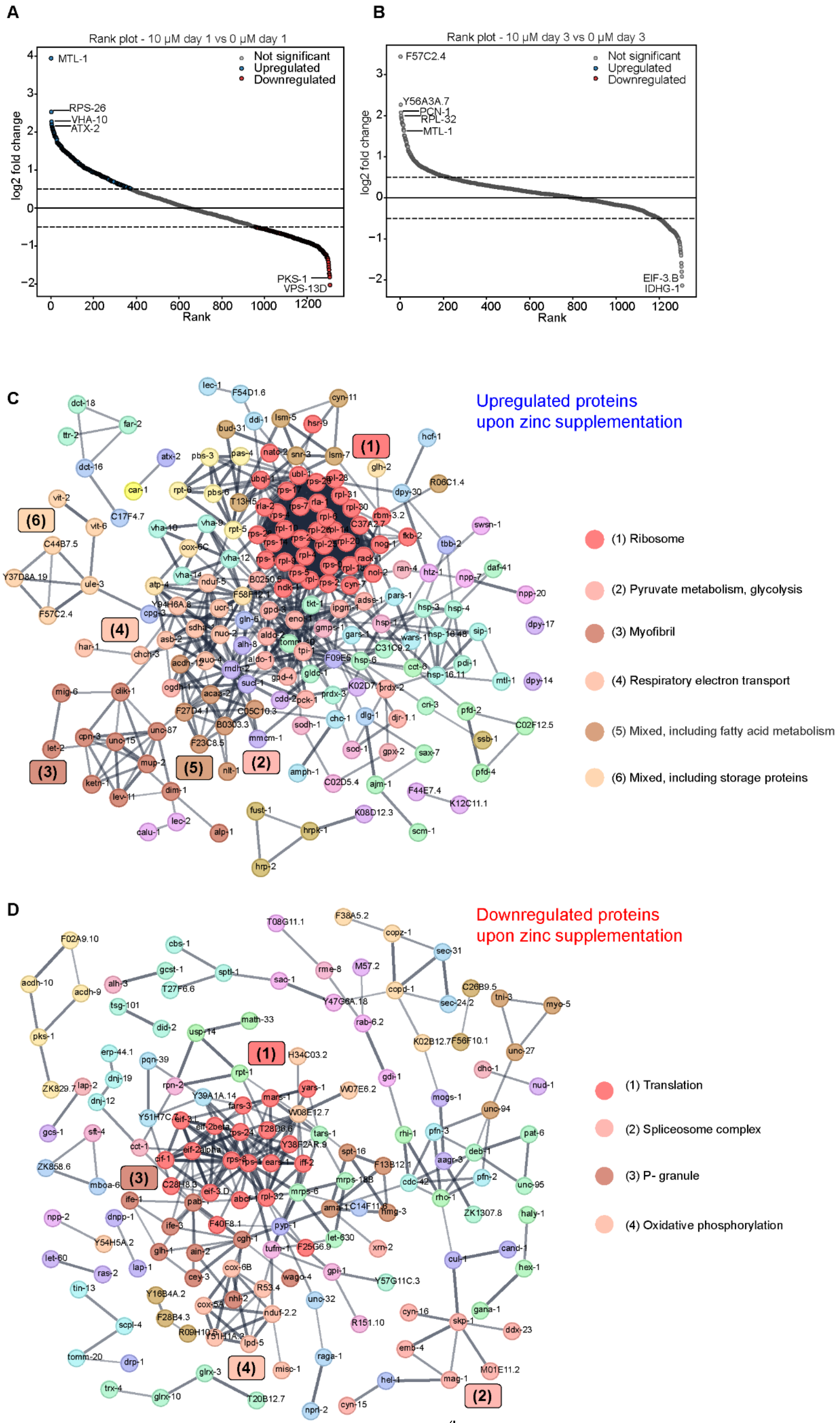
A
Rank plot - 10 µM day 1 vs 0 µM day 1
Not significant
Upregulated
Downregulated
MTL-1
RPS-26
VHA-10
ATX-2
PKS-1
VPS-13D
log2 fold change
Rank
B
Rank plot - 10 µM day 3 vs 0 µM day 3
F57C2.4
Y56A3A.7
PCN-1
RPL-32
MTL-1
EIF-3.B
IDHG-1
C
Upregulated proteins
upon zinc supplementation
(1) Ribosome
(2) Pyruvate metabolism, glycolysis
(3) Myofibril
(4) Respiratory electron transport
(5) Mixed, including fatty acid metabolism
(6) Mixed, including storage proteins
D
Downregulated proteins
upon zinc supplementation
(1) Translation
(2) Spliceosome complex
(3) P- granule
(4) Oxidative phosphorylation

**Supplementary Figure 2. Rank plots and STRING protein-protein interaction network analysis of the zinc-responsive proteome.** (A, B) Rank plots of protein abundance changes after 10 μM treatment at day 1 (A) and day 3 (B). Proteins were ranked from highest to lowest log2 fold change shown on x-axis ("Rank") and y-axis (log2 fold change). Each dot represents one protein. Proteins significantly upregulated (log2 fold change > 0.5, FDR ≤ 0.05) are highlighted in blue (n=236 for day1; n=0 for day 3), and downregulated (log2 fold change < -0.5, FDR ≤ 0.05) in red (n=249 for day1; n=0 for day 3). Dashed lines are fold-change thresholds. Labelled are proteins with the highest and lowest fold changes and MTL-1. Statistical analysis was performed in Perseus and rank plots were generated in Python 3.10. (C) STRING network of proteins significantly upregulated by 10 μM versus 0 μM $ZnCl_2$ at day 1 of treatment (log2 fold change > 0.5, FDR ≤ 0.05), clustered using the Markov Cluster (MCL) algorithm (Cytoscape 3.7.0, STRING app, inflation parameter 3). Clusters are numbered by size/connectivity: (1) Ribosome; (2) Pyruvate metabolism, glycolysis; (3) Myofibril; (4) Respiratory electron transport; (5) Mixed, including fatty acid metabolism; (6) Mixed, including storage proteins. (D) As in (C), for proteins significantly downregulated by zinc supplementation (log2 fold change < -0.5, FDR ≤ 0.05). Clusters: (1) Translation; (2) Spliceosome complex; (3) P granule; (4) Oxidative phosphorylation.

**Supplementary Table 3**

**RT-qPCR primers used in this study**

| Name | Sequence (5'→3') | Purpose | Reference |
|---|---|---|---|
| *cdc-42 Forward* | CTGCTGGACAGGAAGATTACG | RT-qPCR | This study |
| *cdc-42 Reverse* | CTCGGACATTCTCGAATGAAG | RT-qPCR | This study |
| *mtl-1 Forward* | ATGGCTTGCAAGTGTGACTG | RT-qPCR | This study |
| *mtl-1 Reverse* | CACATTTGTCTCCGCACTTG | RT-qPCR | This study |
| *sod-1 Forward* | GCGATTTTTCCGCAGGTCG | RT-qPCR | This study |
| *sod-1 Reverse* | TGGTCATTTTCGGACTTCTG | RT-qPCR | This study |
| *prdx-2 Forward* | GAATTGCTTTCCGTGGACTC | RT-qPCR | This study |
| *prdx-2 Reverse* | AACTGGAAGGCCTGAACAAG | RT-qPCR | This study |
| *gcs-1 Forward* | AAGGTAGTCCGTTGACGTGG | RT-qPCR | This study |
| *gcs-1 Reverse* | TGAGCAATTCCGTGCTTCTTG | RT-qPCR | This study |
| *hsp-6 Forward* | GGAGCTCCAGCCAAAAGAC | RT-qPCR | This study |
| *hsp-6 Reverse* | GGAACGACCTTTAAGTCC | RT-qPCR | This study |
| *pfd-2 Forward* | GCAGAATAATATCGCTAATTTG | RT-qPCR | This study |
| *pfd-2 Reverse* | CGGCAGATTCTTTCTCAG | RT-qPCR | This study |
| *ucr-1 Forward* | GGGGAAAACTTGAGAGAATGGC | RT-qPCR | This study |
| *ucr-1 Reverse* | AGCATTGCGAACCTGGCTAT | RT-qPCR | This study |
| *atp-4 Forward* | GCTCAACAGATTGGCCACCA | RT-qPCR | This study |

| Name | Sequence (5'→3') | Purpose | Reference |
|---|---|---|---|
| *atp-4 Reverse* | GAACTGCAGAGTCAACCTTAGC | RT-qPCR | This study |
| *rpt-5 Forward* | TGTATGGACCACCAGGAACTG | RT-qPCR | This study |
| *rpt-5 Reverse* | CTGAACAAGTTGTGGGCCTG | RT-qPCR | This study |
| *pbs-3 Forward* | CAAGCCACAAGTGTTGTCCG | RT-qPCR | This study |
| *pbs-3 Reverse* | GTGTCGTCCAAACCAGCAAC | RT-qPCR | This study |
| *ubql-1 Forward* | GTCACACCGTCCACTTGGTT | RT-qPCR | This study |
| *ubql-1 Reverse* | CATTCCTCCCATTGTTGGTGC | RT-qPCR | This study |
| *rpl-4 Forward* | CTTCATCGCCGATCAGGTCC | RT-qPCR | This study |
| *rpl-4 Reverse* | GAGTGTTGCTTTCCGGCCT | RT-qPCR | This study |
| *rps-26 Forward* | GCACATCGGAATCCACCCTT | RT-qPCR | This study |
| *rps-26 Reverse* | TGGTGATAGCGACCTGACAGT | RT-qPCR | This study |
| *atg-18 Forward* | TTGGTCCGAAGTCAGACACT | RT-qPCR | This study |
| *atg-18 Reverse* | TGGTCTAAACGGATATGCTTGCT | RT-qPCR | This study |
| *lgg-1 Forward* | ACCCAGACCGTATTCCAGTG | RT-qPCR | [1] |
| *lgg-1 Reverse* | ACGAAGTTGGATGCGTTTTC | RT-qPCR | |
| *snb-1 Forward* | GAATCATGAAGGTGAACGTGG | RT-qPCR (pre-rRNA processing) | [2] |
| *snb-1 Reverse* | CCAATACTTGCGCTTCAGGG | RT-qPCR (pre-rRNA processing) | |
| *#1 pre-rRNA Forward* | CTGTGTTTACACCCGAATGATTCTAG | RT-qPCR (pre-rRNA processing) | |

| Name | Sequence (5'→3') | Purpose | Reference |
|---|---|---|---|
| *#1 pre-rRNA Reverse* | CTAATCGTGAGATGGGACACTCATACA | RT-qPCR (pre-rRNA processing) | |
| *#2 pre-rRNA Forward* | CGCAGACATATAGTCTAGCGAG | RT-qPCR (pre-rRNA processing) | |
| *#2 pre-rRNA Reverse* | GATCCATAGATATTGCTGATGATTC | RT-qPCR (pre-rRNA processing) | |
